# Integrated Tick Management in the Upper Midwest: Impact of Invasive Vegetation Removal and Host-targeted Acaricides on *Ixodes scapularis* Infestation and *Borrelia burgdorferi* Prevalence of Small Mammals

**DOI:** 10.1101/2022.01.07.475376

**Authors:** Jordan T. Mandli, Xia Lee, Susan M. Paskewitz

**Author notes:** Susan Paskewitz, University of Wisconsin-Madison, 1630 Linden Dr. Madison, WI 53706.

## Abstract

Integration of tick management strategies has been suggested to overcome ecological variation in tick, host, pathogen, and habitat, yet empirical evidence assessing combined treatment effect on blacklegged ticks, *Ixodes scapularis* Say, is limited. In this 5-year study (2014-2018) we tested whether combining two methods targeting tick/mammal interactions could reduce juvenile *I*. scapularis parasitism of two small mammal species, *Peromyscus leucopus* Rafinesque and *Tamias striatus* Linnaeus. Infection of small mammals with *Borrelia burgdorferi* was used to evaluate host exposure to feeding ticks. Using a factorial design, removal of invasive vegetation (Amur honeysuckle, *Lonicera maackii* Ruprecht and common buckthorn, *Rhamnus cathartica* Linnaeus) was coupled with deployments of permethrin-treated cotton nesting materials (tick tubes) and evaluated against control sites. Removal of invasive vegetation resulted in lower captures of *T. striatus* suggesting that treatment impacted reservoir activity in the plots. Deployments of permethrin-treated cotton were effective at reducing the frequency of juvenile *I. scapularis* parasitism of *P. leucopus* by 91% across the study compared to controls. However, tick tubes did not offer consistent protection against mouse exposure to *B. burgdorferi* exposure. An additive negative effect was detected for juvenile tick intensity on *P. leucopus* when tick tubes were combined with invasive vegetation removal. We conclude that integration of these two methods provides very limited benefit and that permethrin treatment alone offers the best option for reducing *I. scapularis* infestation on *P. leucopus*.

## 1. INTRODUCTION

Lyme disease (LD) is the most common vector-borne disease in North America and Europe (Bacon et al. 2008). The incidence of LD has steadily increased over the past two decades and the geographic range has expanded dramatically (Kilpatrick et al. 2017). In the northeastern and upper midwestern United States, LD is caused by the spirochete bacteria *Borrelia burgdorferi* sensu stricto and *B. mayonii* (Burgdorfer et al. 1982, Pritt et al. 2016), which are transmitted through the bite of an infected *Ixodes scapularis* Say (blacklegged tick) (Kilpatrick et al. 2017). Blacklegged ticks are also vectors of several other tick-borne pathogens in humans including *B. miyamotoi, Anaplasma phagocytophilum, Babesia microti*, Powassan virus, and *Ehrlichia muris* (Barbour 2017). Continued emergence of tick-borne pathogens necessitates more effective tick control strategies. The complex ecological relationships between habitat, ticks, wild hosts, and tick-borne pathogen often limit broad effectiveness of single interventions (Williams et al. 2018). Integration of more than one of the currently available strategies has been suggested as one way to overcome ecological variation, but empirical evidence for the effectiveness of integrated tick management strategies (ITM) is limited (Bloemer et al. 1990, Williams et al. 2018, Eisen and Stafford 2020).

Juvenile *I. scapularis* ticks will blood feed on a wide variety of vertebrate hosts but the largest proportion of ticks are fed by small mammals (Giardina et al. 2011, Kilpatrick et al. 2017) Vertebrate hosts have varying degrees of reservoir competency, thereby limiting pathogen transmission (Donahue et al. 1987, Telford et al. 1988, Mather et al. 1989). White-footed mice (*Peromyscus leucopus* Rafinesque) are particularly effective at infecting local tick populations with *B. burgdorferi* due to their abundance and extended infectivity (Linske, Williams, Stafford, et al. 2018) and juvenile ticks parasitizing these hosts are highly likely to successfully feed to completion. These mice are also adept at colonizing a variety of habitats, including fragmented tracts in residential neighborhoods, and establishing high population densities when conditions permit (Barko et al. 2003, Nupp and Swihart 2011). When they are in close proximity to humans and domestic animals, *P. leucopus* sustain and infect ticks thereby increasing the risk of local exposure to tick-borne disease. In addition to mice, other hosts such as shrews (*Blarina brevicauda* Say) and chipmunks (*Tamias striatus* Linnaeus) are known to be highly spirochetemic reservoirs perpetuating the *B. burgdorferi* enzootic cycle (Schmidt and Ostfeld 2001, LoGiudice et al. 2003, Brisson et al. 2008)

One method of tick control that targets the host is the commercially available “tick tube”. Tick tubes offer a permethrin-treated nesting material which white-footed mice gather and use towards the construction of their nests. Superficial application of the acaricide to mice kills parasitizing ticks and prevents pathogen transmission (Spielman 1988). Studies have revealed reduced infestation of larval ticks on mice as well as a decrease in the abundance of host-seeking nymphs in the year following tick tube deployment (Mather et al. 1987, 1988, Deblinger and Rimmer 1991). However, small-scale residential studies in New York and Connecticut found that use of tick tubes caused a significant reduction in mouse infestation by ticks but did not reduce questing nymphs following treatment (Daniels et al. 1991, Stafford 1991, 1992, Ginsberg 1992). Variability of mouse treatment has been documented and remains a potential area of improvement; however, has never been extensively evaluated.

Habitat modification has also been used for tick control, especially in relation to invasive plant species. Invasives are characterized by rapid infiltration of the local ecosystem and establishment as the dominant species thereby threatening native biological diversity (Mattos and Orrock 2010). While biological invasions have a direct effect on competing species, the indirect consequences on wildlife communities have been shown to influence ticks and pathogen transmission (Allan et al. 2010). These exotic plants may facilitate interaction between ticks and competent hosts in two ways; 1) modification of the density and distribution of reservoirs by offering cover from predators and/or providing an ample food source for generalist feeders (Berryman and Hawkins 2006, Allan et al. 2010, Mattos and Orrock 2010, Linske, Williams, Ward, et al. 2018), or 2) generation of favorable microclimates that modify tick desiccation-induced tick mortality and allow sustained periods of unhindered questing that increase the probability of tick encounter with hosts (Stafford 1994, Williams et al. 2009). Dense-growing invasive species such as Japanese barberry (*Berberis thunbergii* de Candolle), common buckthorn (*Rhamnus cathartica* Linnaeus) and Amur honeysuckle (*Lonicera maackii* Ruprecht) have been positively associated with questing tick abundance (Lubelczyk et al. 2004, Elias et al. 2006). Removal of invasive plants can reduce the abundance of *B. burgdorferi* infected *I. scapularis* adults that are seeking hosts as well as juvenile ticks that are parasitizing *P. leucopus;* this is thought to occur because of modification of abiotic conditions in altered stands (Williams et al. 2009, 2017, Williams and Ward 2010, Linske, Williams, Ward, et al. 2018). However, the generalizability of these results to other invasive plant species requires further study.

We reported that implementation of invasive vegetation removal or permethrin-treated cotton reduced the density of host-seeking *B. burgdorferi*-infected *I. scapularis* nymphs across a multi-year study (Mandli et al. 2021). However, when these interventions were combined as a part of an integrated approach no additive effect was observed. To further elucidate the underlying mechanisms of these interventions, we examined points of hosts and vector contact. These included treatment effect on tick infestation prevalence, intensity of infestation, and *B. burgdorferi* prevalence in two small mammal species, *P. leucopus* and *T. striatus*. Evaluation of treatment influence on small mammal abundance and activity was completed through animal captures.

## 2. MATERIALS AND METHODS

### 2.1 Study location

Field work took place at the University of Wisconsin (UW) Arboretum located in Madison, WI (43.047764, -89.422732) between May 2014 and August 2018. All sites were established in a temperate forest dominated by red oak (*Quercus rubra* Linnaeus) with associated shagbark hickory (*Carya ovata* Miller), and white pine (*Pinus strobus* Linnaeus). Dense invasive vegetation encompassed a majority of the forest understory. This location was confirmed as having an established *I. scapularis* population in 2012.

In 2014, sixteen non-adjacent 50 × 50 m experimental sites were established, representing the average size of residential lots based on United States home sale census data from 2013 (U.S. Census Bureau 2017). Site sizes (0.25 ha) approximate the lower end of the estimated home range of the white-footed mouse (0.2-0.6 ha). Distance between sites ranged from 35-610 m indicating some potential for mouse cross plot activity (Wolff 1985). Each site was verified for the growth of common buckthorn (*Rhamnus cathartica*) and Amur honeysuckle (*Lonicera maackii)*, while hosting comparable concentrations of native plant species. Prior to the 2015 field season, an additional control site was created and one of the original sites was relocated due to flooding concerns.

### 2.2 Experimental design

Sites were randomly assigned to one of four treatment combinations consisting of vegetation removal and permethrin-treated cotton in a two-by-two factorial design. Treatments consisted of the following: *control* sites underwent no invasive vegetation removal and received tick tubes containing untreated cotton nest material (CTRL), *vegetation removal-only* sites underwent invasive vegetation removal and received tick tubes containing untreated cotton nest material (VR), *permethrin-treated cotton-only* sites underwent no invasive vegetation removal and received tick tubes containing permethrin-treated cotton nest materials (PTC), and *combined* sites received both invasive vegetation removal and tick tubes containing permethrin-treated cotton nesting material (PTC + VR).

### 2.3 Invasive vegetation removal

During plot establishment in May 2014 and 2015 (relocation of site and establishment of new site), common buckthorn and Amur honeysuckle were cleared using loppers, handsaws, and brush cutters from a 20 × 20 m central plot embedded in the full 50 × 50 m experimental site. Non-target plant species and fallen trees were left undisturbed. In years following plot establishment, new growth of the targeted invasive species was removed prior to each field season in early May. An impact of vegetation removal on infesting *I. scapularis* immature ticks and the prevalence of *B. burgdorferi* infection in small mammals is expected within the same year of treatment. Assuming VR treatment influences small mammal activity in plots, captures are expected to decrease immediately.

### 2.4 Tick tube deployment

Twenty-five tubes were deployed across each site approximately 10 m apart, in line with recommended densities of commercial tick tubes and with an emphasis on placement at preferred rodent habitat at the base of trees or along downed logs. Permethrin-treated cotton was prepared from the Tengard SFR One Shot commercial permethrin product during 2014-2016 (United Phosphorus Inc., King of Prussia PA, USA) or the ProZap Insectrin X 10% permethrin concentrate in 2017 and 2018 (CHEM-TECH, LTD., Lexington KY, USA) (providers were changed due to availability). Stock solutions were diluted with water to produce a 10% experimental formulation that was applied to cotton using a 7.6 liter pump sprayer (Roundup®, Marysville, OH). Untreated cotton was prepared with water in a similar fashion, but in a separate location using dedicated equipment. Cotton tube reservoirs were prepared by cutting 6.1 cm (2.4 in) x 3.05 m (10 ft) PVC tubing into lengths of approximately 15.2 cm (6.0 in). Cotton was stuffed into each tube until a 2.54 cm (1 in) lip without cotton was generated at both ends (17 cotton balls/tube). Tick tube deployment occurred once prior to the peak of nymphal and larval activity during 2014-2015 (mid-May) and twice in 2016-2017 (mid-May and early to mid-July) with the second deployment designed to target ticks later in the season. Tubes were recovered from the field in October or November prior to snowfall, cleaned, and reused each year. An impact of permethrin-treated cotton on infesting *I. scapularis* ticks and the prevalence of *B. burgdorferi* infection in white-footed mice should be detectible within the same year of treatment. PTC is expected to have no impact on small mammal captures.

Cotton nesting material uptake was monitored in 2014 through 2018 at the time of second tick tube deployment or tube collection. A lack of observed uptake in early 2017 and 2018 prompted the quantification of material following the first deployment (May-July: ∼ 55 days later) and again following the second deployment (July-October: ∼ 70 days later) in both years.

### 2.5 Small mammal capture

Animal trapping and sampling procedures were reviewed and approved by the Institutional Animal Care and Use Committee of the University of Wisconsin-Madison (#A005400). Small mammals were trapped using Sherman live traps (H.B. Sherman Traps Inc., Tallahassee, FL, USA) once a month from June to August for three consecutive days. A pre-bait night was included to acclimate mice to the presence of traps and increase trapping success. Traps were initially baited with steel-cut oats and an apple chuck as a source of water, however, low animal captures in June 2014 prompted the addition of peanut butter to the bait formulation thereafter. Traps were checked three times a day at 8:00, 12:00, and 16:00 to capture a broad spectrum of nocturnal and diurnal species; white-footed mice, eastern chipmunk, eastern grey squirrel (*Sciurus carolinensis* Gmelin), southern flying squirrel (*Glaucomys volans* Linnaeus), and northern short-tailed shrew. Each site contained five pairs of Sherman traps deployed in a cross-like orientation; a single pair was located at the center of the site and four pairs were placed 10 m away in each cardinal direction (Figure 1). Traps were placed in areas of increased small mammal activity and away from tick tubes.

**Figure 1:**
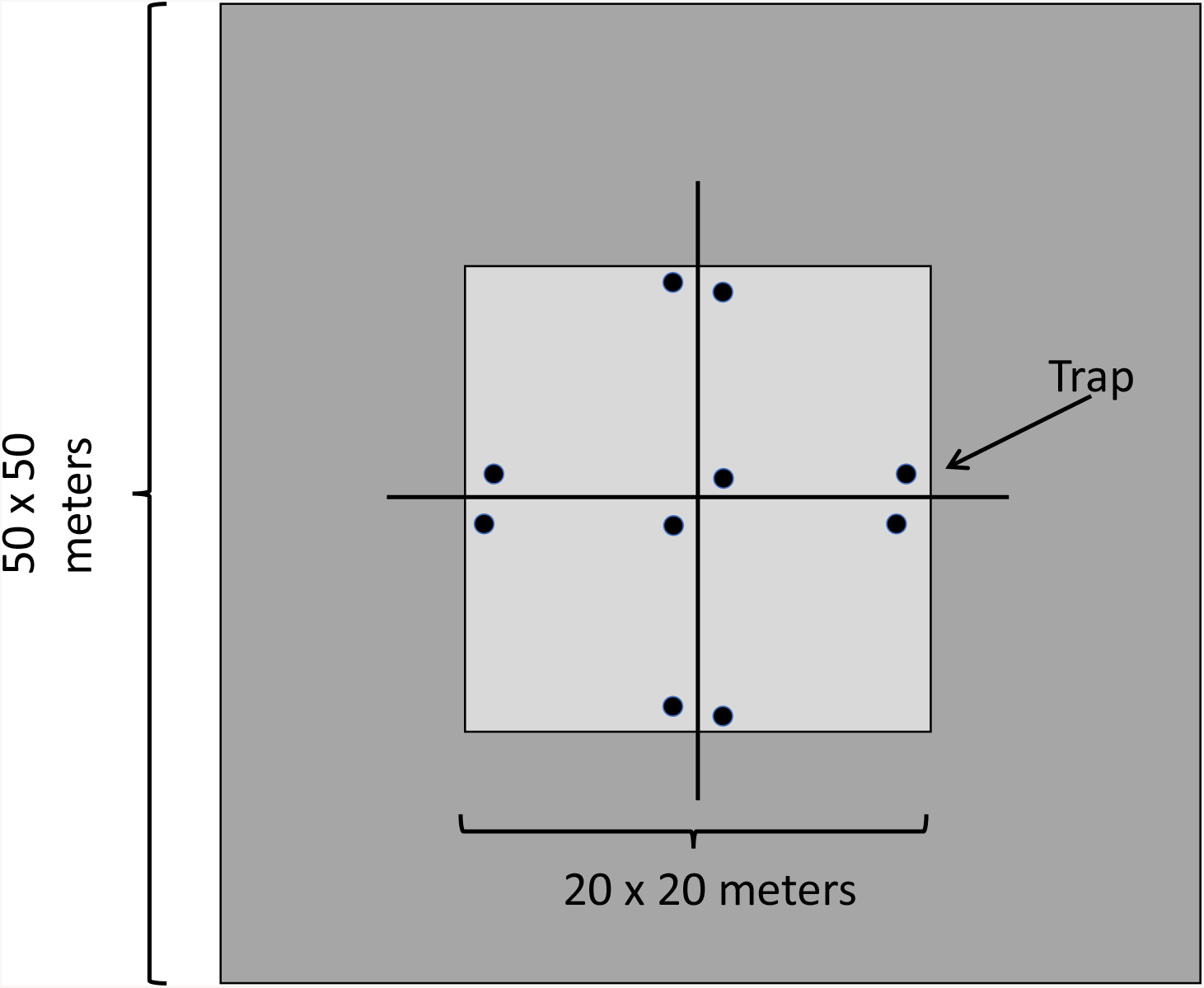
Trapping schematic and dimensions for individual plot. Traps are represented by black circles.

Animal processing occurred onsite. Captured small mammals were identified by species, aged, sexed, and weighed. Small mammals were temporarily sedated with isoflurane, fitted with a uniquely numbered ear tag, and a 2.0 mm ear biopsy was collected and stored in 70% ethanol. Ticks removed from animals were placed in 70% ethanol until identification. All ticks were identified to species and life stage using established taxonomic keys (Clifford et al. 1961, Durden and Keirans 1996). If animals were recaptured within the same month, they were checked for ticks and freed without further processing. Once recovered from the effects of isoflurane, animals were released at the origin of capture.

### 2.6 B. burgdorferi *s*.*s. screening of ear biopsies*

Genomic DNA was extracted from bisected ear biospies using the Bioline Isolate II Genomic DNA Extraction Kit (Bioline USA Inc., Taunton MA, USA) or the DNeasy Blood and Tissue Kit (Qiagen, Valencia, CA) in accordance with the manufacturer’s animal tissue protocol. Samples were tested for *B. burgdorferi* s.s DNA by nested polymerase chain reaction (PCR) targeting the outer surface protein B operon as previously described (Caporale and Kocher 1994, Lee et al. 2014).

### 2.7 Statistical Analysis

#### Nesting material uptake

For 2017 and 2018, the number of remaining cotton balls were summarized as mean residual cotton balls per tick tube by treatment by site during each deployment. Differences between PTC and untreated cotton (UTC) removal were compared using Welch’s two sample t-tests.

The effect of the treatments, PTC, VR and PTC+VR, on animal captures, juvenile *I. scapularis* infestation prevalence, intensity of infestation, and host infection prevalence for two species (*P. leucopus* and *T. striatus*) was assessed through multivariate analyses using generalized linear models (GLMs) for individual years and generalized linear mixed effect models (GLMMs) when accounting for random effects in the cumulative study. Data from recaptured animals in a given trapping session (year + month) were condensed to a single data point (distinct animal) per plot for data analysis. The cumulative 5-year model fixed effects included *Year, Month, Treatment. Site* was included as a random effect in GLMMs to control for natural variation among plots and autocorrelation (Richer et al. 2014, Cayol et al. 2018). Annual GLMs included treatments and month as fixed effects. Model selection was carried out based on corrected Akaike Information Criterion for small sample sizes (AICc) (Burnham and Anderson 2002). The simplest model containing treatment effects within 2 AICc of the model with the lowest AICc value was selected. Post-hoc tests assessing relative excess risk due to interaction (RERI-delta method) and multiplicative interaction were completed (VanderWeele and Knol 2014). Analysis was carried out in R, version 1.4.1717 (R Core Team 2021). GLMMs and GLMs were developed in the lme4 package (Bates et al. 2015). Analyses utilized CTRL as reference unless otherwise stated.

#### Small mammal captures (SMC)

All capture events regardless of recapture status, summarized as the number of *P. leucopus* or *T. striatus* captures per trap day by plot x trapping session. Captures were modeled using negative binomial regression (with log-link function) and relative risk was expressed as incidence rate ratios (IRRs) with a 95% confidence interval.

#### Tick infestation prevalence (INF)

Reported as a positive bivariate outcome (i.e. the presence or absence of at least one immature tick on a host). Direct comparison of the prevalence of infestation by tick life stage between *P. leucopus* and *T. striatus* was completed using a chi-square analysis with Yates’ continuity correction. IP was modeled using logistic regression (with logit-link distribution) and relative risk was expressed as odds ratios (OR) with a 95% confidence interval.

#### Tick intensity of infestation (INT)

Density of immature ticks (larvae and nymphs) parasitizing all infested individual hosts. BI was modeled using negative binomial regression (with log-link function) and relative risk was expressed as IRRs with a 95% confidence interval.

#### B. burgdorferi *infection in hosts (BbP)*

Reported as a positive bivariate outcome (ie: infected vs uninfected host). Direct comparison of infection prevalence between *P. leucopus* and *T. striatus* was completed using a chi-square analysis with Yates’ continuity correction. BbP was modeled using logistic regression (with logit-link distribution) and relative risk was expressed as ORs with a 95% confidence interval.

## 3. RESULTS

### 3.1 Treatment implementation

Initial removal of invasive vegetation was completed in both 2014 and 2015, but continued maintenance was required in order to keep Amur honeysuckle and common buckthorn from reinvading sites. Cotton was actively removed from tick tubes during the first three years (2014 -2016) with less than 5% of cotton nesting material remaining in tubes or on the ground nearby. In 2017, 48% (3456/7225) of the cotton balls remained at the conclusion of the first deployment (May-July). Permethrin-treated cotton was removed less than UTC with a mean 11.2 (95% CI: 9.53 – 12.8) and 5.44 (3.59 – 7.38) residual balls per tube respectively (t = 4.18, p-value < 0.001). During the second deployment in 2017 (July-October), only 3% (227/7225) of the cotton nesting material was left uncollected. At the second deployment, no difference in removal of treated or untreated nest material was detected with an average 0.695 (95% CI: 0.390 – 1.04) (PTC) and 0.391 (95% CI: 0.187– 0.618) (UTC) residual balls per tube (t = 1.399, p-value = 0.19). In 2018, twenty-two tick tubes could not be found following the first deployment. Of those recovered, 27% (1865/6851) of the cotton balls remained. Similar to 2017, PTC was removed less frequently than UTC with a mean 8.13 (6.29 – 9.80) and 1.46 (0.64 – 2.57) balls remaining per tube respectively (t = -6.09, p-value < 0.001). Values following the second deployment were poorly catalogued, limiting our ability to quantitatively evaluate cotton uptake. Still, anecdotal observations indicate near complete cotton removal of both cotton treatments by the end of the 2018 season, matching what was reported in previous years.

### 3.2 Small rodent capture

Over the course of this study, we sampled small mammals on 7,560 trap days; 16 sites were trapped on 3 occasions (1440 trap days/year) in 2014 and 17 sites were trapped on 3 occasions (1530 trap days/year) from 2015-2018. Ten traps were consistently placed at each site while trapping. During our study only two small rodent species were reliably captured and maintained tick burdens, *P. leucopus* (0.181 captures/trap day) and *T. striatus* (0.060 captures/trap day). We encountered other mammal species including *G. volans* (0.002 captures/trap day), *B. brevicauda* (0.001 captures/trap day), and *S. carolinensis* (0.001 captures/trap day). Overall, 901 *P. leucopus* were captured 1,371 times and 266 *T. striatus* were captured 456 times (Suppl. Table 1 and 2). Rates of recapture varied between mammal species; white-footed mice were captured one to six times (601, 180, 81, 30, 7, 2 mice respectively) and chipmunks were captured one to seven times (161, 59, 28, 8, 5, 3, 1 chipmunk respectively) with a single outlier that was captured 11 times. Because some plots were relatively close together, mice and chipmunks were sometimes recaptured on another plot. Of the 901 unique mice that were trapped, 2.2% (20) were also captured on another treatment site at some time. Of the 266 chipmunks that were captured, 4.5% (12) were recaptured at another treatment site. This suggests that movement between plots was low.

The likelihood of *P. leucopus* capture was not significantly different between treatments at the annual and cumulative 5-year level with the exception of PTC in 2015 (Tables 1A & 2). Captures exhibited clear temporal variation in which the likelihood of *P. leucopus* capture increased in July and August compared to June (Table 1A). Using 2014 as reference, the probability of white-footed mouse capture significantly declined during 2015 and 2017 but recovered to similar levels in 2016 and 2018 thereby suggesting a fluctuating two-year trend in the population. The likelihood of *T. striatus* capture was not associated with treatments across the entire study period; however, captures were negatively associated with VR in 2014, 2015, and 2016 (Tables 1B & 3). Similar to mice, chipmunk captures exhibited temporal variability. Across the study, the probability of capturing a chipmunk was significantly lower in August compared to June. Using 2014 as reference, the likelihood of *T. striatus* capture significantly increased in 2017 and 2018 by 80% and 126% respectively.

**Table 1.**
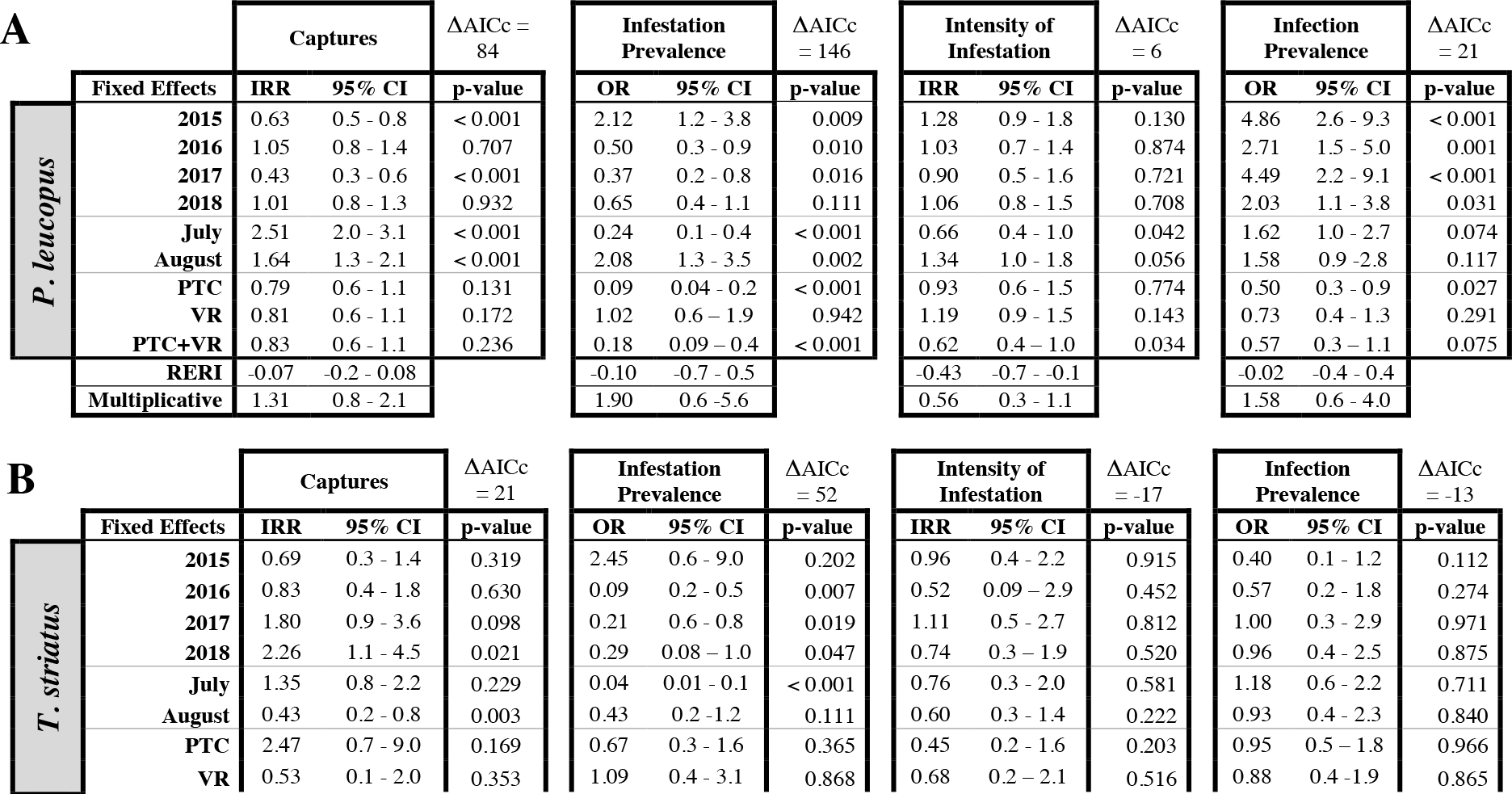
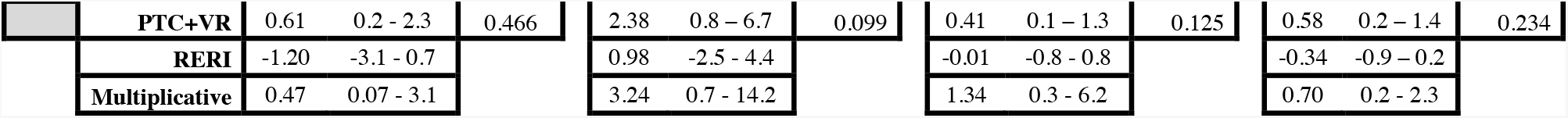
Results from the impact of year, month, permethrin-treated cotton (PTC), vegetation removal (VR), and the combination of PTC and VR on small mammal captures (SMC), juvenile tick infestation prevalence (INF), tick intensity of infestation (INT), *B. burgdorferi* infection prevalence (BbP) across the study for two small mammal species: A) *Peromyscus leucopus* and B) *Tamias striatus* (Data from GLMM). Baseline effects are year: 2014, month: June, Treatment: CTRL. Additive (RERI) and multiplicative interaction terms reported between treatments Differences between minimal model and best model given as **Δ** AICc.

**Table 2.**
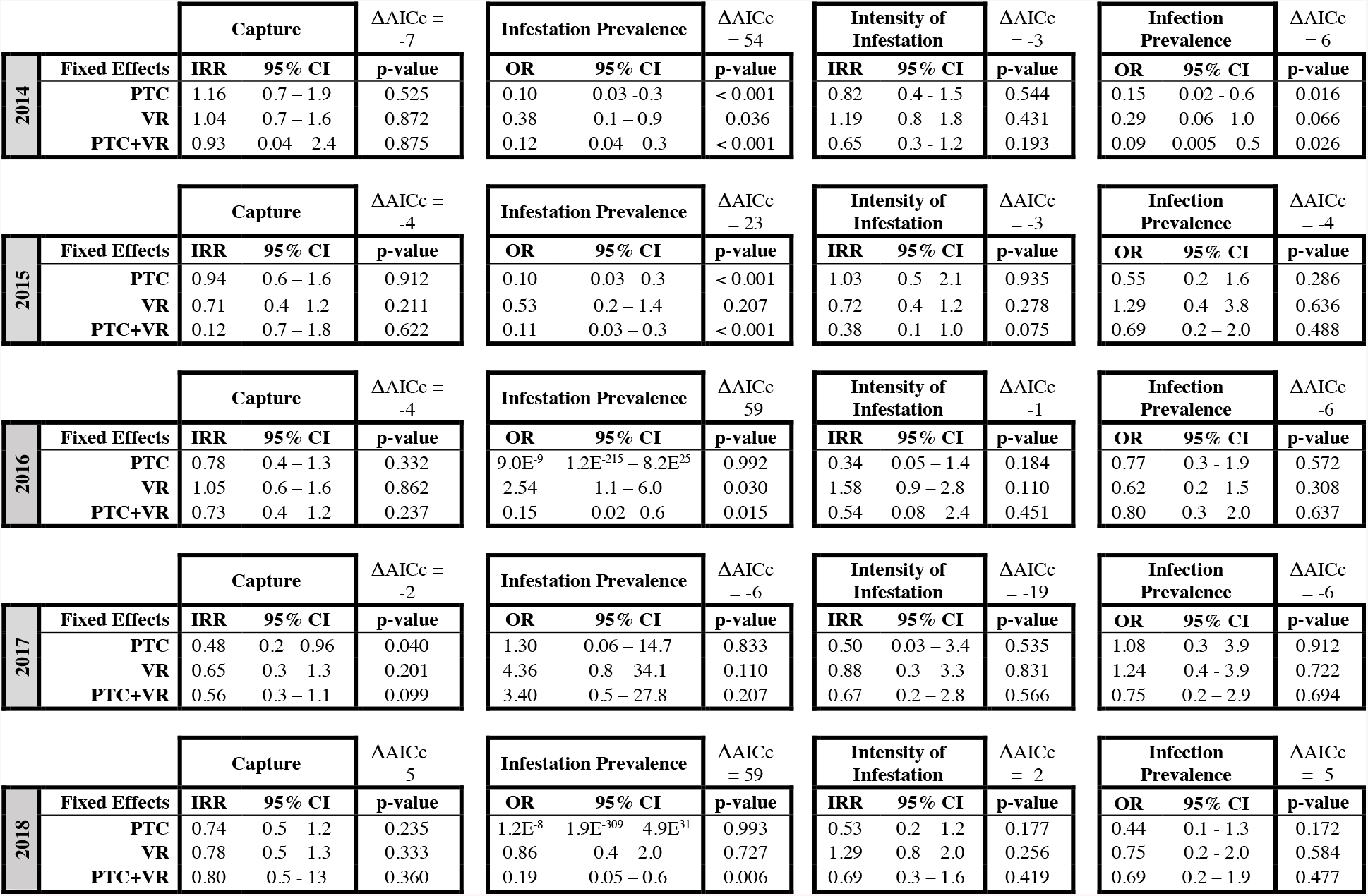
Results from the impact of permethrin-treated cotton (PTC), vegetation removal (VR), and the combination of PTC and VR on small mammal captures (SMC), juvenile tick infestation prevalence (INF) and intensity of infestation (INT), *B. burgdorferi* infection prevalence (BbP) by each year for ***P. leucopus*** (Data from GLM). Baseline effect is Treatment: CTRL. Differences between minimal model and best model given as **Δ** AICc.

**Table 3.** Results from the impact of permethrin-treated cotton (PTC), vegetation removal (VR), and the combinaiton of PTC and VR on small mammal captures (SMC), juvenile tick infestation prevalence (INF) and intensity of infestation (INT), *B. burgdorferi* infection prevalence (BbP) by each year for ***T. striatus*** (Data from GLM). Baseline effect is Treatment: CTRL. Differences between minimal model and best model given as **Δ** AICc.

| | Fixed Effects | Capture | | $\Delta AICc = -1$ | Fixed Effects | Infestation Prevalence | | $\Delta AICc = -2$ | Fixed Effects | Intensity of Infestation | | $\Delta AICc = -11$ | Fixed Effects | Infection Prevalence | | $\Delta AICc = -8$ |
| --- | --- | --- | --- | --- | --- | --- | --- | --- | --- | --- | --- | --- | --- | --- | --- | --- |
|  |  | OR | 95% CI | p-value |  | OR | 95% CI | p-value |  | IRR | 95% CI | p-value |  | OR | 95% CI | p-value |
| 2014 | PTC | 0.62 | 0.2 – 2.5 | 0.498 | PTC | 5.36 | 0.9 – 4.5 | 0.078 | PTC | 1.20 | 0.3 – 8.2 | 0.823 | PTC | 0.33 | 0.03 – 2.6 | 0.309 |
|  | VR | 0.16 | 0.03 – 0.8 | 0.024 | VR | 5.6E<sup>-8</sup> | N/A – 2.8E<sup>241</sup> | 0.996 | VR | 1.67 | 0.4 – 5.8 | 0.429 | VR | 0.66 | 0.06 – 7.5 | 0.733 |
|  | PTC+VR | 0.29 | 0.07 – 1.2 | 0.086 | PTC+VR | 3.00 | 0.2 – 45.5 | 0.395 | PTC+VR | 2.00 | 0.4 – 14.4 | 0.423 | PTC+VR | 0.66 | 0.08 – 5.3 | 0.697 |
| | Fixed Effects | Capture | | $\Delta AICc > 5$ | Fixed Effects | Infestation Prevalence | | $\Delta AICc = -3$ | Fixed Effects | Intensity of Infestation | | $\Delta AICc = -5$ | Fixed Effects | Infection Prevalence | | $\Delta AICc = -6$ |
|  |  | OR | 95% CI | p-value |  | OR | 95% CI | p-value |  | IRR | 95% CI | p-value |  | OR | 95% CI | p-value |
| 2015 | PTC | 0.56 | 0.1 – 2.5 | 0.411 | PTC | 0.42 | 0.1 – 1.5 | 0.187 | PTC | 0.68 | 0.3 – 1.5 | 0.343 | PTC | 0.71 | 0.2 – 3.0 | 0.633 |
|  | VR | 0.13 | 0.02 – 0.6 | 0.005 | VR | 0.56 | 0.07 – 3.6 | 0.549 | VR | 1.23 | 0.4 – 3.1 | 0.682 | VR | 1.22 | 0.1 – 9.6 | 0.848 |
|  | PTC+VR | N/A | N/A | N/A | PTC+VR | N/A | N/A | N/A | PTC+VR | N/A | N/A | N/A | PTC+VR | N/A | N/A | N/A |
| | Fixed Effects | Capture | | $\Delta AICc > 2.0$ | Fixed Effects | Infestation Prevalence | | $\Delta AICc = N/A$ | Fixed Effects | Intensity of Infestation | | $\Delta AICc = N/A$ | Fixed Effects | Infection Prevalence | | $\Delta AICc = -7$ |
|  |  | OR | 95% CI | p-value |  | OR | 95% CI | p-value |  | IRR | 95% CI | p-value |  | OR | 95% CI | p-value |
| 2016 | PTC | 1.65 | 0.5 – 6.0 | 0.411 | PTC | N/A | N/A | N/A | PTC | N/A | N/A | N/A | PTC | 0.67 | 0.1 – 3.8 | 0.646 |
|  | VR | 0.07 | 0.003 – 0.5 | 0.020 | VR | N/A | N/A | N/A | VR | N/A | N/A | N/A | VR | 6.4E<sup>7</sup> | 0.0 – N/A | 0.996 |
|  | PTC+VR | 0.59 | 0.1 – 2.4 | 0.453 | PTC+VR | N/A | N/A | N/A | PTC+VR | N/A | N/A | N/A | PTC+VR | 1.00 | 0.1 – 9.1 | 0.999 |
| | Fixed Effects | Capture | | $\Delta AICc = -1$ | Fixed Effects | Infestation Prevalence | | $\Delta AICc = 1$ | Fixed Effects | Intensity of Infestation | | $\Delta AICc = -15$ | Fixed Effects | Infection Prevalence | | $\Delta AICc = -4$ |
|  |  | OR | 95% CI | p-value |  | OR | 95% CI | p-value |  | IRR | 95% CI | p-value |  | OR | 95% CI | p-value |
| 2017 | PTC | 2.68 | 0.8 – 9.7 | 0.118 | PTC | 0.07 | 0.004 – 0.5 | 0.023 | PTC | 0.15 | 0.006 – 3.2 | 0.187 | PTC | 2.33 | 0.6 – 9.4 | 0.220 |
|  | VR | 1.47 | 0.4 – 5.4 | 0.546 | VR | 0.26 | 0.03 – 1.4 | 0.139 | VR | 0.45 | 0.08 – 3.1 | 0.375 | VR | 0.70 | 0.2 – 3.1 | 0.638 |
|  | PTC+VR | 0.45 | 0.1 – 1.8 | 0.245 | PTC+VR | 1.92 | 0.1 – 50.8 | 0.643 | PTC+VR | 0.15 | 0.01 – 1.28 | 0.075 | PTC+VR | 0.58 | 0.08 – 7.7 | 0.689 |
| | Fixed Effects | Capture | | $\Delta AICc = -6$ | Fixed Effects | Infestation Prevalence | | $\Delta AICc = -3$ | Fixed Effects | Intensity of Infestation | | $\Delta AICc = -11$ | Fixed Effects | Infection Prevalence | | $\Delta AICc = -5$ |
|  |  | OR | 95% CI | p-value |  | OR | 95% CI | p-value |  | IRR | 95% CI | p-value |  | OR | 95% CI | p-value |
| 2018 | PTC | 1.87 | 0.6 – 6.5 | 0.312 | PTC | 2.25 | 0.3 – 47.0 | 0.494 | PTC | 1.33 | 0.3 – 9.6 | 0.740 | PTC | 0.83 | 0.3 – 2.7 | 0.763 |
|  | VR | 1.21 | 0.4 – 4.3 | 0.766 | VR | 6.32 | 0.9 – 126.5 | 0.105 | VR | 2.40 | 0.7 – 15.4 | 0.252 | VR | 1.19 | 0.3 – 4.4 | 0.789 |
|  | PTC+VR | 1.24 | 0.4 – 4.4 | 0.727 | PTC+VR | 2.08 | 0.3 – 20.0 | 0.478 | PTC+VR | 1.33 | 0.3 – 9.6 | 0.740 | PTC+VR | 0.45 | 0.1 – 1.6 | 0.235 |

### 3.3 *Juvenile* Ixodes scapularis *parasitizing small rodents*

A total of 530 juvenile ticks (478 larvae, 52 nymphs) were collected from all small rodents; 414 juvenile ticks (403 larvae / 11 nymphs: CTRL = 6, VR = 2, PTC = 0, PTC+VR = 3) were obtained from *P. leucopus* and 116 juvenile ticks (75 larvae / 41 nymphs) from *T. striatus*. Larvae were more likely to infest mice (OR = 1.70, *χ*^2^ = 6.29, df 1, p-value = 0.012) while nymphs were more likely to infest chipmunks (OR = 5.82, *χ*^2^ = 16.9, df 1, p-value < 0.001). No other captured mammal species presented any observed *I. scapularis*. Across the study, 17% (n = 1101) of all distinct white-footed mice were infested with at least one juvenile tick, whereas 16% (n = 318) of all distinct chipmunks were infested. Mean intensity of infestation was 2.27 (median: 2.0, range: 1-13) ticks per distinct *P. leucopus* and 2.27 (median: 1.0, range: 1-26) ticks per distinct *T. striatus*.

The prevalence of infestation exhibited substantial temporal variation that coincided with the tick phenology discussed in Mandli et al. 2021 (Mandli et al. 2021); *P. leucopus* infestation corresponded to larval questing patterns and *T. striatus* infestation matched peak nymphal patterns (Table 1A&B).

As for treatments, PTC significantly reduced the likelihood of infestation on white-footed mice by 91% across the study but had no significant impact on INT (Table 1A). PTC was negatively associated with IP in 2014 and 2015 and exhibited complete inhibition of infestation on mice in 2016 and 2018 (Table 2). In 2017, CTRL exhibited a large decline in infestation prevalence which impacted comparative assessments of PTC. Even so, mice captured on PTC sites exhibited nearly complete absence of tick infestation. At the cumulative level, VR exhibited no significant impact on the probability of infestation. However, during the first year of treatment *P. leucopus* INF was negatively associated with VR, with a 62% decrease. The combination of PTC and VR was negatively associated with INT at the cumulative level. As such, additive interaction was detected between treatments for *P. leucopus* (Table 1A). The likelihood of tick infestation on *T. striatus* was not associated with treatments at the annual or cumulative 5-year level (Table 1B & 3). Moreover, none of the explored effects were associated with INT in chipmunks.

### 3.4 *Small rodent* B. burgdorferi *infection*

A strong dichotomy existed between chipmunk and mouse *B. burgdorferi* infection (OR = 5.15, *χ*^2^ = 115.8, df 1, p-value < 0.001). In total 14% (142/1013) of *P. leucopus* and 46% (105/230) of *T. striatus* ear biopsies from unique animals were infected with *B. burgdorferi*. Thirteen mice became infected with *B. burgdorferi* over the course of a given season; three were captured in CTRL plots, four from VR plots, four from PTC plots, and two from PTC+VR plots. Of note, one of these mice moved from a CTRL to a PTC+VR plot and another moved from CTRL to PTC so they may have become infected outside PTC-treated plots. Annual *B. burgdorferi* prevalence in *P. leucopus* and *T. striatus* was variable (Table 1A&B). The likelihood of *B. burgdorferi* infection in *P. leucopus* was negatively associated with PTC (= lower infection) at the cumulative level but remained variable in each individual year (Table 1A). Infection with *B. burgdorferi* in chipmunks was not associated with any explored effects.

## 4. DISCUSSION

Results from this study demonstrate that PTC is a reliable deterrent of juvenile blacklegged tick parasitism on *P. leucopus*. This affirms the causal mechanism of PTC reduction of DON and DIN as effective inhibition of tick and host contact. PTC resulted in immediate reductions to INF on white-footed mice that persisted throughout much of the study; however, intensity of infestation on infested mice did not differ from CTRLs suggesting an all or nothing response. Reductions to tick infestation imposed by VR were short-lived. Significant reductions to INF on mice were constrained to the first year of study; however, intensity of infestation did not differ from CTRL. Despite reductions in INF for both treatments in 2014, no interaction was detected. Neither treatment exhibited an impact on *T. striatus* tick infestation or intensity of infestation. This was not unexpected; chipmunks exhibit little interest in PTC and have a larger home range which takes them well outside the boundaries of the VR plots.

Permethrin-treated nesting materials are expected to inhibit all tick feeding, including infected nymphs, in an attempt to disrupt spirochete transmission. As such, we expected that *P. leucopus* captured in sites with deployed tick tubes would have lower *B. burgdorferi* infection compared to CTRL sites. While PTC exhibited a cumulative reduction to prevalence of *B. burgdorferi* in mice, the impact was inconsistent from year to year, probably because the prevalence of infection and the number of mice tested per treatment each year were low. Invasive vegetation removal had no influence on *B. burgdorferi* prevalence in mice. Given a lack of effect on tick parasitism, it was no surprise that treatments had no impact on chipmunk infection with *B. burgdorferi*.

Dense vegetation provides refuge for small mammals, especially white-footed mice, thereby facilitating interaction between ticks and hosts (Allan et al. 2010, Mattos and Orrock 2010, Guiden and Orrock 2017). As such, we initially hypothesized that removal of invasive vegetation would decrease mouse relative abundance. However, this study showed that the likelihood of capturing a *P. leucopus* was indistinguishable between treatments annually and across the cumulative 5-year study. Linske et al (2018b) reported a similar lack of difference in *P. leucopus* captures after the removal of Japanese barberry from 30 × 30 m plots. Surprisingly, *T. striatus* exhibited lower rates of capture among VR plots in 2014-2016, perhaps suggesting that removal of invasive plants may act as a deterrent to activity of this diurnal species. PTC does not offer any advantage in small mammal survivability (Mather et al. 1987); therefore, it was no surprise that chipmunk and mouse captures in the PTC plots were not different from CTRLs. While not the main focus of this study, our data shows significant variation in *T. striatus* and *P. leucopus* captures between years as well as among months. This is likely a result of fluctuating environmental and ecological factors including weather (Madison et al. 1984), predator abundance (Mattos and Orrock 2010), and food availability (Ostfeld, Jones, et al. 1996) and population dynamics (Wang et al. 2008), and seasonal population growth (Wolff 1986, Mather et al. 1987).

Results from this study suggest that permethrin-treated cotton may offer complete protection against *I. scapularis* infestation in an area of low tick abundance. Across the study, only 6% of all distinct mice captured (n = 490) from PTC or PTC+VR sites were infested with at least one tick. The body burdens on these infested mice were similar compared to CTRLs suggesting that these may be mice that originated from untreated locations and indeed, we documented movement of mice from control to treated plots. Other PTC trials have similarly documented all or nothing responses (Mather et al. 1987, Deblinger and Rimmer 1991). Furthermore, Ginsberg et al. (1992) demonstrated that mice with detectable levels of permethrin on hair were completely free of parasitizing ticks. Assuming PTC treatment of mice offers complete protection against infestation, animals with parasitizing ticks in areas of permethrin treatment are likely untreated. Exposing mouse movement patterns under different conditions as well as nesting behaviors related to acaricide application are key to refining this strategy for broader use.

We saw an overall reduction in the prevalence of *B. burgdorferi* in mice captured in PTC plots, but sampling was underpowered and low prevalence and small sample sizes precluded the ability to detect annual patterns for most years. Reducing host infection is key to minimizing human risk because individual mice can infect hundreds of ticks. Given that the action of permethrin-treated cotton includes the inhibition of infected nymph feeding on mice, we expected that *P. leucopus* captured in PTC and PTC+VR sites would have lower *B. burgdorferi* infection compared to CTRL sites. Our data revealed a complete lack of parasitizing nymphs at PTC sites and three instances of nymph parasitism at PTC+VR sites. While frequency of parasitizing nymphs was low, a few mice captured in sites with permethrin-treated cotton became infected over the season. However, several of these mice originated in untreated areas. Studies describing the impact of PTC on the prevalence of *B. burgdorferi* in mice are lacking, but other host-targeted systems, like fipronil-bait boxes, have described an inability to reduce prevalence of infection of *P. leucopus* at small scales (< 1.0 ha) (Dolan et al. 2004, Schulze et al. 2017). Both host-targeted approaches are constrained by an area of effect. Assuming mice closer to a tick tube have a higher probability of becoming treated, a larger area of treatment is more likely to generate protection at its center. Mouse home ranges are estimated to be approximately 0.2-0.6 hectares (0.5-1.5 acres), but this can vary depending on population density and season (Wolff 1985). Landscape features (man-made structures), other nesting substrates, and mice with unusually expansive home ranges could further complicate estimates of area of effect. Further investigation is needed to understand spatial constraints of PTC and other host-targeted approaches in order to refine deployment strategies.

Spirochete infection in mice may also occur prior to PTC treatment. Tick tubes were deployed at the same time each year, however increasingly warmer temperatures earlier in the season may have advanced tick phenologies and increased mouse and tick interactions prior to treatment (Levi et al. 2015). Another possible explanation is the reported presence of immature ticks in mouse nests. Larson et al. 2019 found unfed blacklegged tick nymphs among nests collected in May. Moreover, ticks were also observed in nests collected in August, suggesting the possibility of overwintering in these locations. Given our lack of understanding of the longevity of permethrin treatment on mice and the stability of the acaracide on cotton substrates in the field, we do not know if PTC in nests remains effective by spring. Therefore, it is possible that mouse exposure begins in the nest. Further research assessing the longevity and transferability of permethrin in the field is needed to determine when deployments are most efficacious. If nymphal tick overwintering in nests or early questing does occur, it may be effective to provide PTC late in the fall prior to mouse torpor.

Low cotton collection in 2017 was likely a function of a mouse abundance. Mouse captures were significantly lower than any other year of study. As a result, *P. leucopus* home ranges may have increased and mice may have moved more outside the bounds of our study sites (Wolff 1985, Ostfeld, Miller, et al. 1996). Mice exhibited lower levels of INF in CTRL sites that resembled PTC sites suggesting cross treatment. All three instances of cross plot migration that we observed in 2017 were a result of mice moving between areas of permethrin-treated cotton to those without. Under these conditions, PTC appeared less effective as compared to CTRL likely as a result of some treated mice entering these areas. The use of multi-year studies (> 2) is necessary for interpretation of these low-density events.

Invasion of exotic vegetation can have a dramatic impact on local ecology thereby modifying host and tick interactions. However, across this study, VR had little impact on capture rates, tick infestation frequency and intensity, and *B. burgdorferi* prevalence of *P. leucopus*, yet significant reductions in DIN (density of infected questing nymphs) were documented in Mandli et al 2021. These results suggest that some alternative factor besides larval tick survivorship on or infection by white-footed mice in VR plots altered DIN the following year (T+1). The presence of alternative hosts could explain the reduction in densities of infected nymphs. *Tamias striatus* and *B. brevicauda* were both captured on our sites and are known to be competent *B. burgdorferi* reservoirs (Schmidt and Ostfeld 2001, LoGiudice et al. 2003, Brisson et al. 2008). The absence of chipmunks in VR sites coupled with their high rates of *B. burgdorferi* infection could account for declines in DIN (T+1). While chipmunks are typically associated with higher rates of nymphal tick attachment (Schmidt et al. 1999), chipmunks do feed larvae and some of these must contribute to the population of infected nymphs. Future work could benefit from nymph bloodmeal analysis to better understand the sources of tick infection (Cadenas et al. 2007). Interventions that discourage competent hosts or encourage incompetent hosts presence have the potential to dilute pathogen presence (Linske et al. 2018). Strategic application of VR as an intervention will require further investigation of how it influences alternative species in order to justify its application.

Here, we evaluated the impact of integrated environmental control strategies on *B. burgdorferi* prevalence and *I. scapularis* parasitism of two small mammal species in south central Wisconsin. Our results demonstrate that PTC reliably reduced mouse infestation by juvenile ticks, however it does not completely eliminate the role of mice in maintaining spirochete transmission. VR removal had no detectable impact on white-footed mouse infestation, infection, or abundance suggesting an effect on tick feeding and/or spirochete transmission in alternative reservoir hosts. Continued examination of treatment impact on host and tick interactions remains a key part of understanding variation in treatment effectiveness. Additional work is needed to refine these strategies and determine which combinations offer the cost-effective option for the general public.

## Acknowledgments

We thank the following individuals for their support of our research efforts: staff at the UW Arboretum, especially ecologist Brad Herrick; K. Bartowitz and J. Orrock for advice on experimental design and assistance in the field; Sam Engle with the UW College of Agricultural and Life Science Statistical Consulting Center; and our undergraduate and graduate students for their help with field work. Funding: Jordan Mandli was supported by the Parasitology and Vector Biology Training Program T32AI007414. This work was supported by the USDA National Institute of Food and Agriculture, Hatch project 0232758 and the Centers for Disease Control and Prevention, Cooperative Agreement Number U01CK000505. Its contents are solely the responsibility of the authors and do not necessarily represent the official views or position of the Centers for Disease Control and Prevention, the Department of Health and Human Services, or the U.S. Government.

## SUPPLEMENTARY

**Supplementary Table 1:** Mean (median, range) captures per plot trap day, infestation prevalence and mean (median, range) intensity infestation of juvenile *I. scapularis* on *P. leucopus*, and *B. burgdorferi* infection prevalence by treatment and year for *P. leucopus*.

|  |  | *P. leucopus* |  |  |  |
| --- | --- | --- | --- | --- | --- |
|  | Treatment | Captures | Infestation Prevalence | Intensity of Infestation | % Infected |
| 2014 | CTRL | 2.3 (1.8, 0-7) | 35% (25/71) | 2.2 (2.0, 1-5) | 16% (11/70) |
|  | VR | 2.1 (1.7, 0-5) | 20% (12/61) | 2.7 (2.0, 1-8) | 5% (3/59) |
|  | PTC | 2.4 (2.0, 0-6) | 8% (6/77) | 1.8 (1.5, 1-3) | 3% (2/74) |
|  | PTC + VR | 2.1 (2.2, 0-5) | 11% (7/65) | 1.4 (1.0, 1-3) | 2% (1/58) |
|  | Annual Total | 2.2 (1.8, 0-7) | 18% (50/274) | 2.2 (2.0, 1-8) | 7% (17/261) |
| 2015 | CTRL | 1.4 (1.0, 0-3) | 49% (24/49) | 2.7 (2.0, 1-9) | 27% (12/45) |
|  | VR | 1.0 (1.0, 0-2) | 37% (11/30) | 1.9 (1.0, 1-5) | 32% (8/25) |
|  | PTC | 1.4 (1.3, 0-3) | 11% (4/37) | 2.8 (2.0, 1-6) | 17% (6/36) |
|  | PTC + VR | 1.6 (1.3, 0-4) | 11% (4/38) | 1.0 (1.0, 1-1) | 20% (7/35) |
|  | Annual Total | 1.3 (1.3, 0-4) | 28% (43/154) | 2.3 (1.0, 1-9) | 23% (33/141) |
| 2016 | CTRL | 2.6 (2.7, 0-5) | 15% (14/96) | 1.9 (1.0, 1-5) | 18% (16/88) |
|  | VR | 2.7 (2.3, 0-7) | 29% (21/73) | 3.0 (2.0, 1-13) | 12% (8/66) |
|  | PTC | 2.0 (2.3, 0-3) | 0% (0/63) | 0.0 (0.0, NA) | 15% (8/55) |
|  | PTC + VR | 1.9 (2.0, 1-4) | 4% (2/55) | 1.0 (1.0, 1-1) | 15% (8/53) |
|  | Annual Total | 2.3 (2.0, 0-7) | 13% (37/287) | 2.5 (1.0, 1-13) | 15% (40/262) |
| 2017 | CTRL | 1.3 (1.3, 0-4) | 4% (2/46) | 2.0 (2.0, 1-3) | 22% (10/45) |
|  | VR | 0.9 (0.7, 0-2) | 17% (4/24) | 1.8 (1.5, 1-3) | 26% (6/23) |
|  | PTC | 0.6 (0.3, 0-2) | 5% (1/19) | 1.0 (1.0, 1-1) | 24% (4/17) |
|  | PTC + VR | 0.8 (0.7, 0-1) | 15% (3/20) | 1.3 (1.0, 1-2) | 18% (3/17) |
|  | Annual Total | 0.9 (0.7, 0-4) | 9% (10/109) | 1.6 (1.0, 1-3) | 23% (23/102) |
| 2018 | CTRL | 2.6 (2.3, 0-5) | 24% (24/101) | 2.2 (1.0, 1-8) | 15% (13/86) |
|  | VR | 2.0 (2.0, 0-4) | 23% (14/60) | 2.9 (2.0, 1-8) | 12% (6/51) |
|  | PTC | 1.9 (1.7, 0-4) | 0% (0/59) | 0.0 (0.0, NA) | 7% (4/55) |
|  | PTC + VR | 2.1 (1.7, 0-6) | 7% (4/57) | 1.5 (1.5, 1-2) | 11% (6/55) |
|  | Annual Total | 2.2 (2.0, 0-6) | 15% (42/277) | 2.3 (2.0, 1-8) | 12% (29/247) |
| Cumulative | CTRL | 2.0 (1.7, 0-7) | 25% (89/363) | 2.3 (2.0, 1-9) | 19% (62/334) |
|  | VR | 1.7 (1.3, 0-7) | 25% (62/248) | 2.6 (2.0, 1-13) | 14% (31/224) |
|  | PTC | 1.7 (1.3, 0-6) | 4% (11/255) | 2.1 (2.0, 1-6) | 10% (24/237) |
|  | PTC + VR | 1.7 (1.3, 0-6) | 9% (20/235) | 1.3 (1.0, 1-3) | 11% (25/218) |
|  | Project Total | 1.8 (1.3, 0-7) | 17% (182/1101) | 2.3 (2.0, 1-13) | 14% (142/1013) |

**Supplementary Table 2:** Mean (median, range) captures per plot trap day, infestation prevalence and mean (median, range) intensity of infestation of juvenile *I. scapularis* on *T. striatus*, and *B. burgdorferi* infection prevalence by treatment and year for *T. striatus*

|  |  | *T. striatus* |  |  |  |
| --- | --- | --- | --- | --- | --- |
|  | Treatment | Captures | Infestation Prevalence | Intensity of infestation | % Infected |
| 2014 | CTRL | 0.8 (0.2, 0-3) | 12% (2/17) | 1.0 (1.0, 1-1) | 60% (6/10) |
|  | VR | 0.1 (0.0, 0-1) | 0% (0/4) | 0.0 (0.0, NA) | 50% (2/4) |
|  | PTC | 0.5 (0.0, 0-3) | 42% (5/12) | 1.2 (1.0, 1-2) | 33% (2/6) |
|  | PTC + VR | 0.2 (0.0, 0-1) | 25% (2/8) | 2.0 (2.0, 1-3) | 50% (3/6) |
|  | Annual Total | 0.4 (0.0, 0-3) | 22% (9/41) | 1.3 (1.0, 1-3) | 50% (13/26) |
| 2015 | CTRL | 0.6 (0.0, 0-4) | 47% (9/19) | 2.4 (1.0, 1-7) | 35% (6/17) |
|  | VR | 0.3 (0.0, 0-1) | 33% (2/6) | 3.0 (3.0, 1-5) | 40% (2/5) |
|  | PTC | 1.0 (0.3, 0-5) | 27% (6/22) | 1.7 (2.0, 1-2) | 28% (5/18) |
|  | PTC + VR | 0.0 (0.0, 0-0) | 0% (0/0) | 0.0 (0.0, NA) | N/A |
|  | Annual Total | 0.5 (0.0, 0-5) | 36% (17/47) | 2.2 (2.0, 1-7) | 33% (13/40) |
| 2016 | CTRL | 0.4 (0.0, 0-2) | 0% (0/13) | 0.0 (0.0, NA) | 33% (3/9) |
|  | VR | 0.03 (0.0, 0-0) | 0% (0/1) | 0.0 (0.0, NA) | 100% (1/1) |
|  | PTC | 0.7 (0.3, 0-3) | 0% (0/18) | 0.0 (0.0, NA) | 31% (4/13) |
|  | PTC + VR | 0.3 (0.0, 0-2) | 38% (3/8) | 1.0 (1.0, 1-1) | 40% (2/5) |
|  | Annual Total | 0.4 (0.0, 0-3) | 8% (3/40) | 1.0 (1.0, 1-1) | 36% (10/28) |
| 2017 | CTRL | 0.6 (0.0, 0-6) | 28% (5/18) | 6.6 (2.0, 1-26) | 46% (6/13) |
|  | VR | 0.9 (0.0, 0-4) | 9% (2/22) | 3.0 (3.0, 1-5) | 38% (6/16) |
|  | PTC | 1.7 (0.8, 0-7) | 3% (1/36) | 1.0 (1.0, 1-1) | 67% (18/27) |
|  | PTC + VR | 0.3 (0.3, 0-1) | 22% (2/9) | 1.0 (1.0, 1-1) | 33% (1/3) |
|  | Annual Total | 0.9 (0.3, 0-7) | 12% (10/85) | 4.2 (1.5, 1-26) | 53% (31/59) |
| 2018 | CTRL | 0.7 (0.2, 0-4) | 4% (1/25) | 1.0 (1.0, 1-1) | 52% (11/21) |
|  | VR | 0.9 (0.2, 0-5) | 21% (5/24) | 2.4 (2.0, 1-4) | 59% (10/17) |
|  | PTC | 1.3 (1.2, 0-3) | 9% (3/35) | 1.3 (1.0, 1-2) | 50% (11/22) |
|  | PTC + VR | 0.9 (0.0, 0-4) | 14% (3/21) | 1.3 (1.0, 1-3) | 35% (6/17) |
|  | Annual Total | 0.9 (0.3, 0-5) | 11% (12/105) | 1.8 (1.5, 1-4) | 49% (38/77) |
| Cumulative | CTRL | 0.6 (0.0, 0-6) | 18% (17/92) | 3.4 (1.0, 1-26) | 46% (32/70) |
|  | VR | 0.4 (0.0, 0-5) | 16% (9/57) | 2.7 (2.0, 1-5) | 49% (21/43) |
|  | PTC | 1.1 (0.7, 0-7) | 12% (15/123) | 1.4 (1.0, 1-2) | 47% (40/86) |
|  | PTC + VR | 0.3 (0.0, 0-4) | 22% (10/46) | 1.3 (1.0, 1-3) | 39% (12/31) |
|  | Project Total | 0.6 (0.0, 0-7) | 16% (51/318) | 2.3 (1.0, 1-26) | 46% (105/230) |

**Supplementary Table 3:** Mean (median, range) captures per plot trap day, infestation prevalence and mean (median, range) intensity of infestation of juvenile *I. scapularis* on *P. leucopus*, and *B. burgdorferi* infection prevalence by treatment and month for *P. leucopus*.

|  |  | *P. leucopus* |  |  |  |
| --- | --- | --- | --- | --- | --- |
|  | Treatment | Captures | Infestation Prevalence | Intensity of infestation | % Infected |
| June | CTRL | 1.3 (1.2, 0-4) | 20% (15/76) | 2.5 (1.0, 1-9) | 13% (9/69) |
|  | VR | 0.8 (0.7, 0-2) | 33% (13/40) | 1.7 (2.0, 1-4) | 12% (4/34) |
|  | PTC | 0.8 (0.3, 0-3) | 11% (4/37) | 2.8 (2.0, 1-6) | 6% (2/33) |
|  | PTC + VR | 1.1 (1.0, 0-3) | 10% (5/48) | 1.2 (1.0, 1-2) | 16% (7/44) |
|  | Monthly Total | 1.0 (0.7, 0-4) | 18% (37/201) | 2.1 (1.0, 1-9) | 12% (22/180) |
| July | CTRL | 2.9 (2.7, 0-7) | 9% (16/171) | 1.3 (1.0, 1-3) | 22% (35/158) |
|  | VR | 2.7 (2.2, 0-6) | 9% (11/126) | 1.5 (1.0, 1-5) | 15% (17/117) |
|  | PTC | 2.5 (2.5, 0-6) | 1% (1/130) | 1.0 (1.0, 1-1) | 10% (12/123) |
|  | PTC + VR | 2.2 (1.8, 0-6) | 4% (4/96) | 1.3 (1.0, 1-2) | 10% (9/88) |
|  | Monthly Total | 2.6 (2.7, 0-7) | 6% (32/523) | 1.3 (1.0, 1-5) | 15% (73/486) |
| August | CTRL | 1.8 (1.5, 0-5) | 50% (58/116) | 2.5 (2.0, 1-8) | 17% (18/107) |
|  | VR | 1.7 (1.0, 0-7) | 46% (38/82) | 3.3 (2.0, 1-13) | 14% (10/73) |
|  | PTC | 1.8 (2.0, 0-5) | 7% (6/88) | 1.8 (1.5, 1-3) | 12% (10/81) |
|  | PTC + VR | 1.7 (1.5, 0-5) | 12% (11/91) | 1.4 (1.1, 1-3) | 10% (9/86) |
|  | Monthly Total | 1.8 (1.3, 0-7) | 30% (113/377) | 2.6 (2.0, 1-13) | 14% (47/347) |

**Supplementary Table 4:** Mean (median, range) captures per plot trap day, infestation prevalence and mean (median, range) intensity of infestation of juvenile *I. scapularis* on *T. striatus*, and *B. burgdorferi* infection prevalence by treatment and month for *T. striatus*.

|  |  | *T. striatus* |  |  |  |
| --- | --- | --- | --- | --- | --- |
|  | Treatment | Captures | Infestation Prevalence | Intensity of infestation | % Infected |
| June | CTRL | 0.8 (0.2, 0-6) | 28% (11/40) | 4.7 (2.0, 1-26) | 45% (14/31) |
|  | VR | 0.6 (0.0, 0-4) | 40% (9/23) | 2.7 (2.0, 1-5) | 50% (8/16) |
|  | PTC | 1.1 (0.3, 0-7) | 21% (8/39) | 1.4 (1.0, 1-2) | 26% (16/28) |
|  | PTC + VR | 0.3 (0.0, 0-2) | 40% (6/15) | 1.2 (1.0, 1-2) | 25% (2/8) |
|  | Monthly Total | 0.7 (0.0, 0-7) | 29 (34/117) | 2.8 (2.0, 1-26) | 45% (37/83) |
| July | CTRL | 0.8 (0.3, 0-4) | 8% (3/36) | 1.0 (1.0, 1-1) | 45% (13/29) |
|  | VR | 0.7 (0.0, 0-5) | 0% (0/30) | 0.0 (0.0, NA) | 46% (11/24) |
|  | PTC | 1.5 (0.8, 0-5) | 2% (1/56) | 2.0 (2.0, 2-2) | 51% (21/41) |
|  | PTC + VR | 0.5 (0.0, 0-4) | 0% (0/19) | 0.0 (0.0, NA) | 40% (6/15) |
|  | Monthly Total | 0.9 (0.3, 0-5) | 3% (4/141) | 1.25 (1.0, 1-2) | 47% (51/109) |
| August | CTRL | 0.3 (0.0, 0-1) | 19% (3/16) | 1.0 (1.0, 1-1) | 50% (5/10) |
|  | VR | 0.1 (0.0, 0-1) | 0% (0/4) | 0.0 (0.0, NA) | 67% (2/3) |
|  | PTC | 0.6 (0.3, 0-3) | 21% (6/28) | 1.3 (1.0, 1-2) | 35% (6/17) |
|  | PTC + VR | 0.2 (0.0, 0-1) | 33% (4/12) | 1.5 (1.0, 1-3) | 50% (4/8) |
|  | Monthly Total | 0.3 (0.0, 0-3) | 22% (13/60) | 1.3 (1.0, 1-3) | 45% (17/38) |

